# LIMD1 phase separation contributes to cellular mechanics and durotaxis by regulating focal adhesion dynamics in response to force

**DOI:** 10.1101/828806

**Authors:** Yuan Wang, Chunlei Zhang, Shipeng Shao, Yujie Sun, Ling Liang, Congying Wu

**Affiliations:** Peking University

## Abstract

The mechanical environement affects cell morphology, differentiation and motility. The ability of cells to follow gradients of extracellular matrix stiffness-durotaxis has been implicated in development, fibrosis, and cancer. Cells sense and respond to extra-cellular mechanical cues through cell-matrix adhesions. Interestingly, the maturation of focal adhesions (FAs) is reciprocally force-dependent. How biomechanical cues dictate the status of cell motility and how FAs coordinate force sensing and self-organization remain enigmatic. LIMD1, a member of the LIM domain proteins, localizes to the FAs and has been reported to negatively regulate the Hippo-YAP pathway in response to tension. Here we identify the force sensitive recruitment of LIMD1 to the FAs. We discover that LIMD1 regulates cell spreading, maintains FA dynamics and cellular force, and is critical for durotaxis. Intriguingly, LIMD1 selectively recruits late but not early FA proteins through phase separation at the FAs under force. We suggest a model in which localization of LIMD1 to the FAs, triggered by mechanical force, serves as a phase separation hub for assembling and organizing late FA proteins, allowing for effective FA maturation and efficient cellular mechano-transduction.

---

Force transmission from the extracellular environment to the interior of a cell has been a fascinating subject for decades. Integrin mediated adhesion sites bind to extracellular molecules and initiate a cascade of molecular assembly underneath the plasma membrane, constructing a highly organized structure that is capable of transducing mechanical force^1^. These adhesion sites have been extensively investigated for their crucial roles in mediating both inside-out and outside-in signaling which are essential for cell survival and motility^1-5^. Focal adhesions (FAs) are cell-matrix contacts that mature from nascent focal complexes containing integrins and a few other molecules such as kindlin, talin, FAK and paxillin^6-8^. Only a portion of the nascent focal complexes survives the maturation process and during that they grow by rapidly recruiting late FA proteins to assemble plaque like structures and clutching to the actin stress fibers. Matured FAs exhibited ordered ultrastructure with the force transmission layer sandwiched between the integrin layer and the actin association layer^9^. Specific protein-protein interactions have been identified to participate in FA maturation^10-12^. However, how the army of FA proteins achieves spatial-temporally controlled recruitment and assembly remains unveiled.

Efficient force transmission through the FAs is essential for numerous cellular processes including cell migration. Durotaxis is a type of directional cell migration in which cells respond to a gradient of extracellular stiffness^13, 14^. Mechanisms underlying durotaxis of mesenchymal cells are proposed to include contractile mechanosensation, probing of the local substrate by actin-based protrusions, and focal adhesion signaling^15-21^. Force fluctuations within individual FAs have been observed to mediate ECM-rigidity sensing in guiding directed cell migration^15^. How the formation and dynamics of the complex protein assemblies at cell-ECM contact sites coordinate with force sensing and cell motility is yet to be thoroughly investigated.

The purification of adhesion complexes combined with quantitative mass spectrometry has enabled the identification and quantification of known and new adhesion-associated proteins^22-24^. Blocking the adhesion maturation with the myosin II inhibitor blebbistatin markedly impaired the recruitment of LIM domain proteins to integrin adhesion sites^24^. LIMD1 is a member of LIM domain-containing proteins and a tumor suppressor^25-27^. Interestingly, LIMD1 has recently been identified to regulate the Hippo/YAP signaling pathway by binding and suppressing LATS1/2 in a tension-dependent manner^28^. The role and mechanism underlying LIMD1 mechano-sensation are unclear.

By immunofluorescent staining, we detected co-localization of LIMD1 and the focal adhesion marker protein paxillin (Fig. 1a). We also noticed that the distribution of LIMD1 fluorescent signal within individual FAs is biased towards the proximal region (towards the cell interior and away from the cell edge), suggestive of its role in FA maturation rather than in early FA formation (Fig. 1a and Supplementary Fig. 1a). Enhanced LIMD1 localization at the focal adhesions was obvious with increased substrate stiffness (Fig. 1b). Similar results were detected when we used blebbistatin to inhibit actomyosin contractility (Fig. 1e). These observations are consistent with previous reports that many LIM proteins of the FA components were recruited in a force-dependent manner^24^. Of note, the total protein level of LIMD1 was indistinguishable between cells on soft or stiff substrates, as well as between cells treated with DMSO or blebbistatin (Fig. 1c and Supplementary Fig. 1b). Interestingly, when cells on soft substrate were stimulated with a RhoA activator (Rho activator II), they recovered FA localization of LIMD1 (Fig. 1d). Meanwhile, when cells on glass were treated with blebbistatin or Y27632 to inhibit actomyosin contraction, they exhibited loss of LIMD1 at the FAs (Fig. 1e). These observations demonstrate sensitivity of LIMD1 localization to substrate stiffness and cellular force. Intriguingly, inhibiting the focal adhesion kinase (FAK) resulted in loss of LIMD1 localization at the FAs even when cells were plated on glass (Supplementary Fig. 1c), suggesting that the force-dependent recruitment of LIMD1 to FAs is under the regulation of early FA signaling events. Collectively, these results claim that the mechanical force, either from substrate stiffness or from cell contractility, acted on recruiting LIMD1 to the FAs.

**Figure 1.**
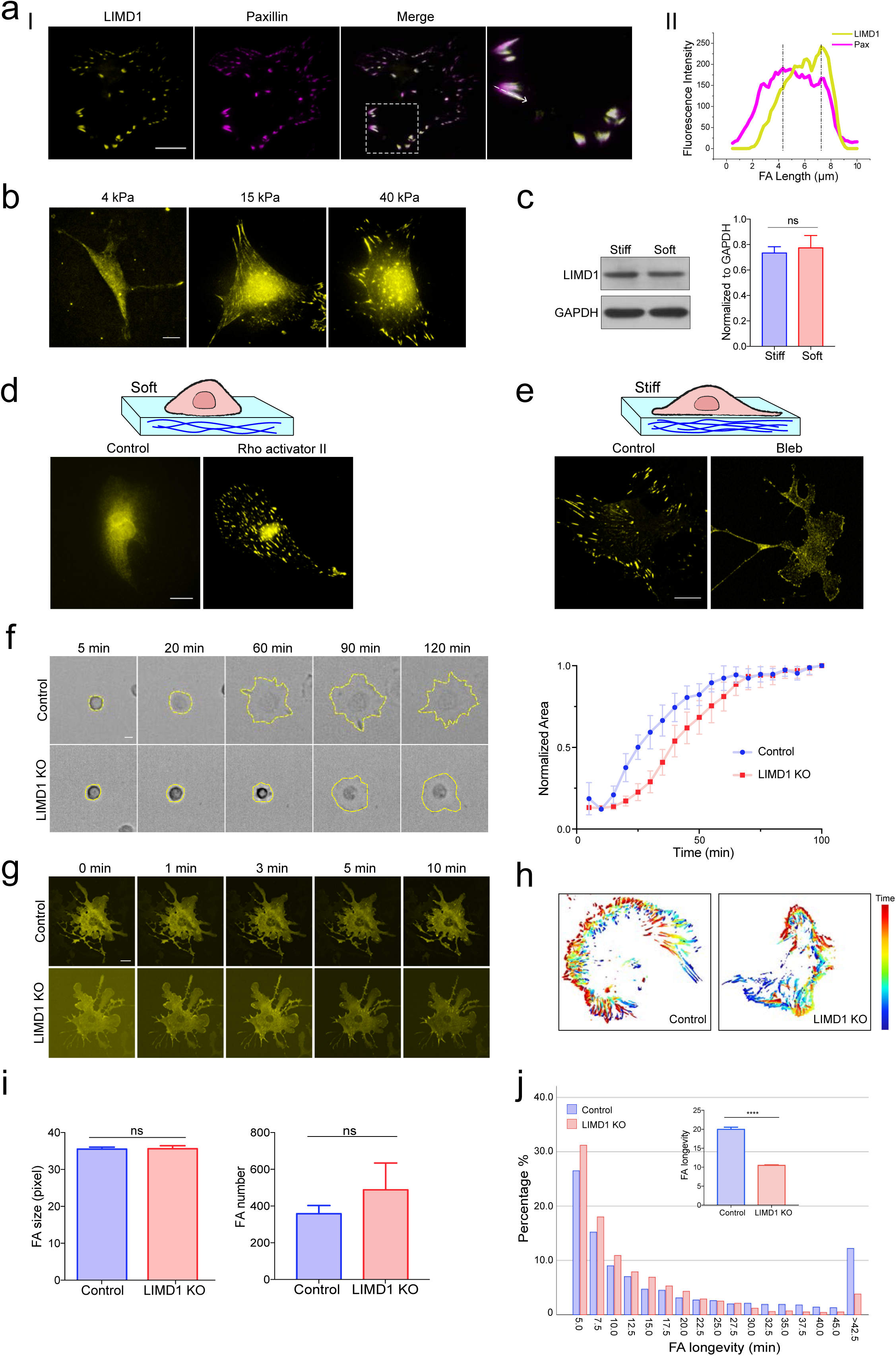
LIMD1 localizes to FAs under force and regulates FA dynamics. **(a)** I: representative immunofluorescence images stained with LIMD1 (yellow) and paxillin (magenta) antibodies in mouse embryonic fibroblasts (MEFs). Localization of LIMD1 and paxillin localization within individual FAs is shown in enlarged boxes. II: Distribution of LIMD1 (yellow) and paxillin (magenta) along the direction of the white arrowhead is shown in the line scan plot. **(b)** Representative immunofluorescence images of LIMD1 localization at FAs on substrates with different stiffness. Scale bars, 20 μm. **(c)** Western blot showing the total protein level of LIMD1 in MEFs on soft and stiff substrates. GAPDH is used as loading control. Bar chart shows quantification of protein levels normalized to GAPDH in each condition (error bars: mean with SEM, ns: not significant). **(d)** Representative immunofluorescence images of LIMD1 on soft substrate with and without 10 μM Rho activator II treatment. Scale bars, 20 μm. **(e)** Representative immunofluorescence images of LIMD1 on stiff substrate with or without 50 μM blebbistatin treatment. Scale bars, 20 μm. **(f)** Left: time-lapse images showing the spreading of control and LIMD1 knockout (LIMD1 KO) MEFs. Scale bar, 20 μm. Right: spreading area curves of control and LIMD1 KO MEFs. Error bar: 95% CI. **(g)** Time-lapse images showing FA formation in control and LIMD1 KO MEFs upon washout of blebbistatin. Scale bar, 20 μm. **(h)** Representative images of FA longevity analyzed by Focal Adhesion Analysis Server (FAAS, faas.bme.unc.edu), color spectrum indicates different time points. **(i)** Bar graphs showing FA size and number (N = 12 cells, error bar: 95% CI, *P < 0.05, ***P < 0.001 by Student’s *t* test for comparison of two groups. ns, not significant). 1 pixel = 0.18333 μm. **(j)** The frequency distribution of FA longevity in control and LIMD1 KO MEFs. The histogram is filtered to only include adhesions with longevity more than 5 min (N =12 cells, error bar: 95% CI, ***P < 0.001 by *Mann Whitney* test).

To probe the cellular function of LIMD1, we generated LIMD1 knockout (LIMD1 KO) cells. We noticed that depletion of LIMD1 delayed cell spreading (Fig. 1f and Supplementary Video 1-2). We thus wondered whether the maturation process of FAs was hampered upon loss of LIMD1. Firstly, we noticed delayed FA formation in LIMD1 KO cells upon washout of blebbistatin (Fig. 1g and Supplementary Video 3-4). Next, to explore the cellular function of LIMD1, we utilized live-cell TIRF and the focal adhesion analysis server (FAAS)^29^ to analyze the FA dynamics in a global manner (Fig. 1h and Supplementary Video 5-6). FA size or number remain unchanged in LIMD1-depleted cells (Fig. 1i). However, we observed significantly decreased FA longevity upon LIMD1 depletion (Fig. 1j). When we plotted individual FA longevity, we noticed a dramatic drop of long-lived FAs in LIMD1 KO cells (Supplementary Fig. 1d). These results reveal an important role of LIMD1 in regulating FA dynamics.

FA maturation is under the regulation of cellular mechanics while it also feeds back to the mechano-transduction and mechanical properties of a cell^30-32^. To interrogate cell contraction, we utilized the gel deformation assay by imbedding control or LIMD1 KO cells in collagen gels. The LIMD1 KO cells failed to deform the gel to the extent as the control cells, indicative of weakened cellular mechanics (Fig. 2a). Moreover, traction force microscopy also revealed reduction in cellular traction exerted by cells lacking LIMD1 (Fig. 2b). Since LIMD1 exhibited force sensitive localization and regulates cellular force, and that FAs are critical structures for mesenchymal cell migration, we then sought to explore whether LIMD1 influences durotaxis--the mechanical cue induced directional cell migration. Polyacrylamide gels with a stiffness gradient of about 30 kPa/mm were manufactured as previously described^17^. Homogenous coating of the extracellular matrix on the gel surface was demonstrated using fluorescently labeled fibronectin (Fig. 2c). This excluded that the directional migration observed was due to any haptotactic cue adhered to the substrate. We recorded live cell movies for both control and LIMD1 KO cells moving on these gels and analyzed the speed, persistence and directionality based on their migration projection. We didn’t notice any significant differences in migration speed or persistence between control and LIMD1 KO cells (Supplementary Fig. 2a,b). However, we found that control cells showed stiff-end biased direction on the gradient gel, while the LIMD1 KO cells still exhibited random motility (Fig. 2d and Supplementary Video 7-8). These results reveal the impairment of cellular mechanics and loss of durotaxis upon depletion of LIMD1 in cells.

**Figure 2.**
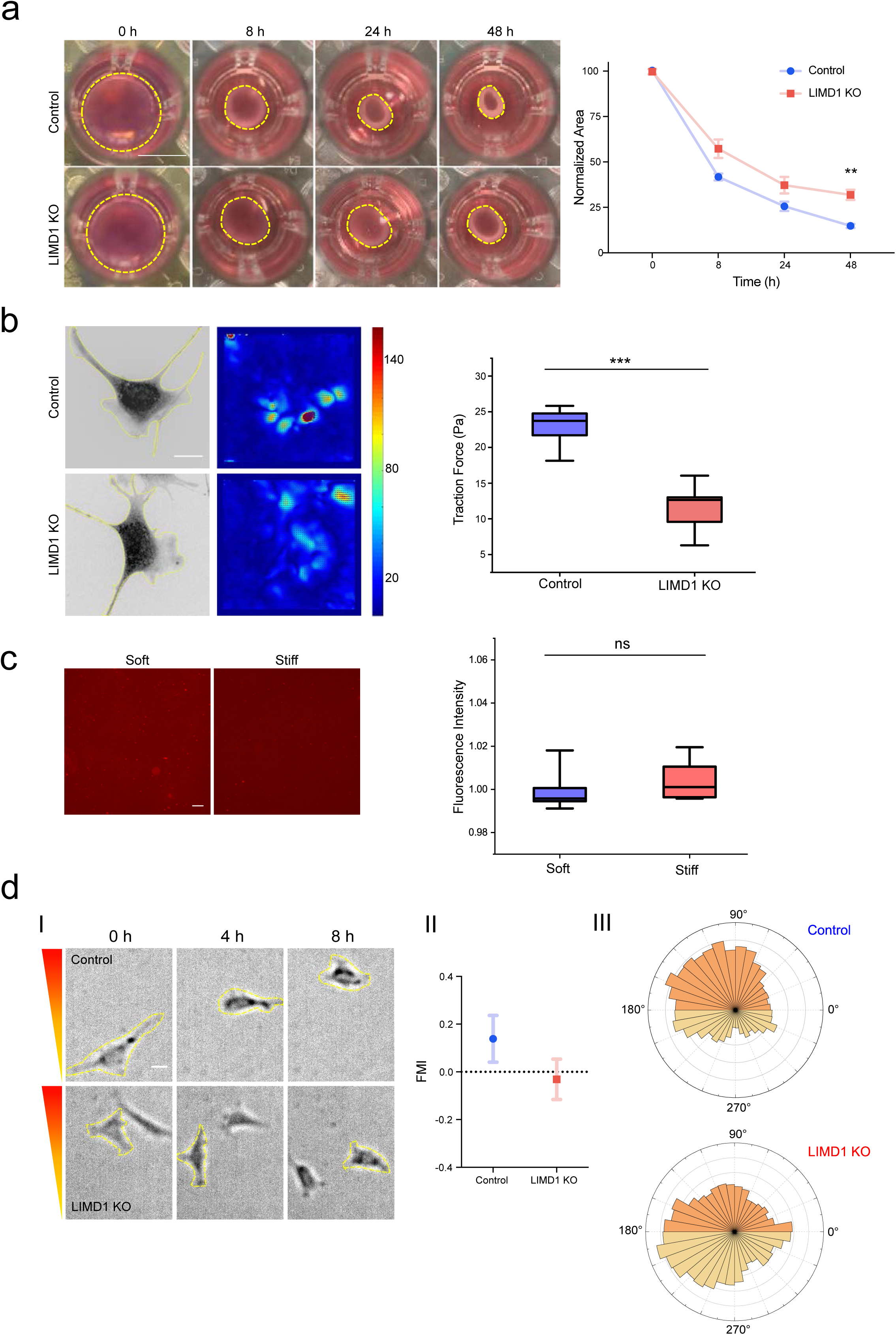
Perturbation of LIMD1 alters cellular mechanics and durotaxis. **(a)** Left: control and LIMD1 KO MEFs are cultured in 3D collagen gels. The morphology of cell-collagen gel mixture is recorded at 0 h, 8 h, 24 h and 48 h. Scale bar, 5 mm. Right: Quantification of collagen gel contraction using measurement of gel area (N = 3 experiments, error bars: mean with SD, *P < 0.05, ***P < 0.001 by Student’s *t* test for comparison of two groups). **(b)** Left: representative images from traction force microscopy (TFM) experiments. Color spectrum indicates stress magnitude (Pa), with areas of low traction in blue and high traction in red. Right: Quantification of total force from TFM experiments showing that LIMD1 KO cells exert less traction force than control cells (N = 5 cells, error bars: 95% CI, *P < 0.05, **P < 0.01, and ***P < 0.001 by Student’s *t* test for comparison of two groups. ns, not significant). Scale bar, 20 μm. **(c)** Right: representative images of Cy3 labeled fibronectin on soft and stiff regions of the stiffness gradient gel. Scale bar, 100 μm. Left: quantification of fibronectin fluorescence intensity of soft and stiff regions (error bars: 95% CI, Student’s *t* test for comparison of two groups. ns, not significant). **(d)** (I): representative time lapse images showing control and LIMD1 KO MEFs during durotactic migration. Scale bar, 20 μm. (II): forward migration index (FMI) of control and LIMD1 KO MEFs. NCon = 43 cells, NKO = 52 cells. (III): rose plots of angle distribution of control (top) and LIMD1 KO (bottom) MEFs as in (II).

LIMD1 contains three LIM domains at the C-terminus with an internal disordered region (IDR) at the N-terminus (Fig. 3a). Using bioinformatics tools, we predicted that the LIMD1 protein has a tendency to undergo phase separation (Fig. 3a). Indeed, when we purified the LIMD1 protein, we observed formation of phase separated droplets in the low salt buffer (Fig. 3b). Dynamic fusion events of these droplets were also captured in the protein containing solution (Fig. 3b and Supplementary Video 9). These droplets showed rapid recovery upon photo-bleaching (Fig. 3c and Supplementary Video 10). In order to examine whether LIMD1 could undergo phase separation in cells, we employed the Opto-droplet based optogenetic system^33^. Exposure to blue light instantaneously induced LIMD1 droplet formation both in the cytoplasm and at the FA sites (Fig. 3d and Supplementary Video 11). In support of LIMD1 undergoing phase separation inside the cells, FRAP experiments showed that the FA retained LIMD1 is dynamic (Fig. 3e and Supplementary Video 12), indicating that the LIMD1 proteins within the FAs rapidly exchanges with their cytoplasmic counterparts. Phase separation has been suggested to rely on multi-valency interactions mediated by IDRs as well as tandemly linked modular domains^34, 35^. We then sought to investigate the critical regions on LIMD1 that contributed to its phase separation property. Surprisingly, truncation of the LIM domains (ΔLIM) or the IDR (ΔIDR) drastically altered the LIMD1 droplet formation kinetics in cells (Fig. 3f and Supplementary Video 13-14). Consistently, loss of the LIM domains significantly deteriorated the phase separation ability of LIMD1 in vitro (Fig. 3g and Supplementary Fig. 3a).

**Figure 3.**
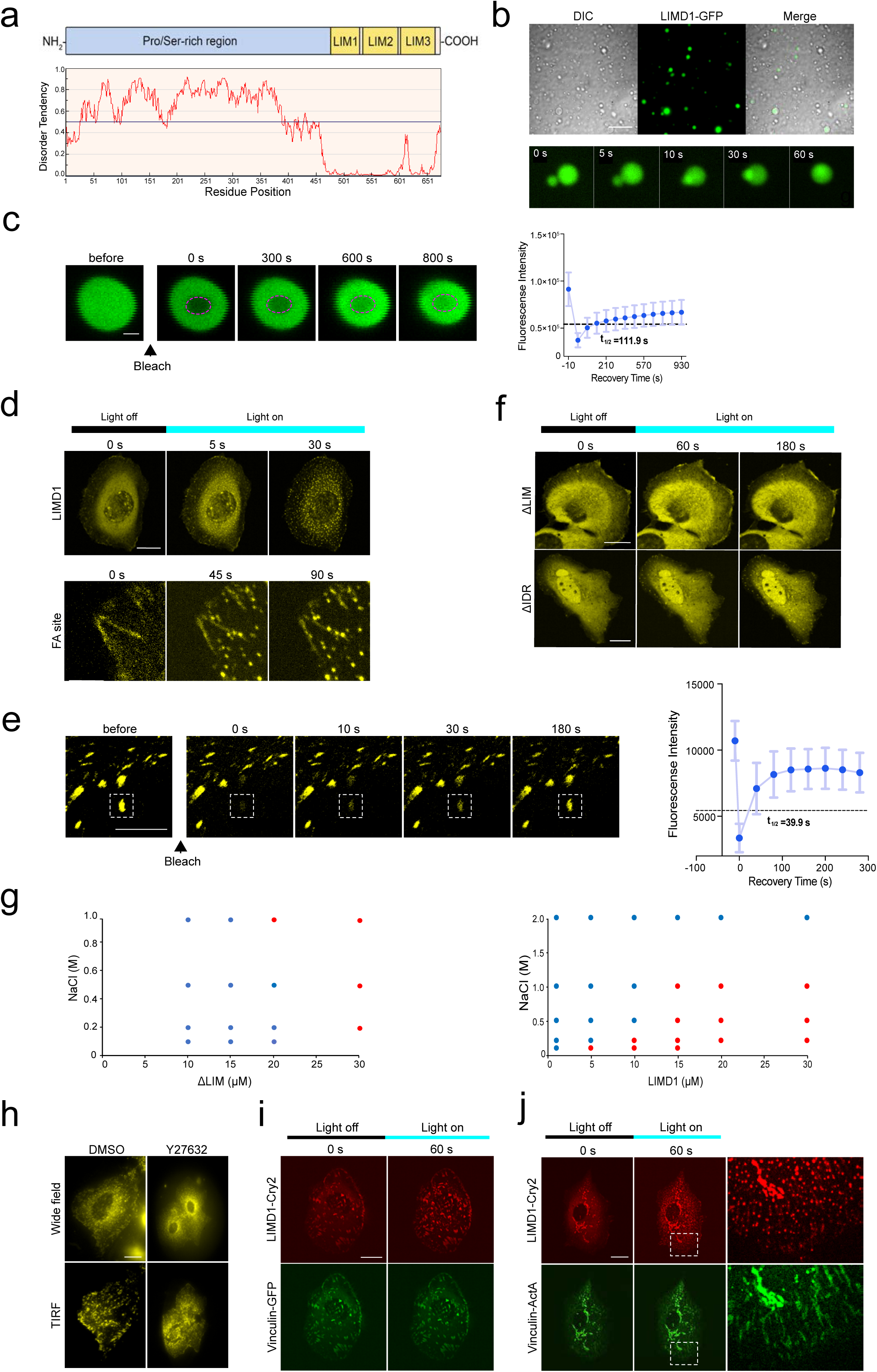
Perturbation of LIMD1 alters cellular mechanics and durotaxis. **(a)** Top: schematic representation of LIMD1 domain structure. Bottom: intrinsic disorder region of LIMD1 predicted by IUPred (http://iupred.elte.hu). **(b)** Top: representative DIC and fluorescence images showing that the LIMD1 protein is concentrated into spherical droplets. Bottom: representative time-lapse images demonstrating small droplets fusing into larger ones. Scale bar, 20 μm. **(c)** FRAP (fluorescence recovery after photobleaching) analysis showing that LIMD1 proteins enriched in the condensed droplets dynamically exchanges with those in the dilute phase. Right: quantification of FRAP analysis of the area indicated by the dashed circle. Scale bar, 2 μm. **(d)** Top: representative images of cells expressing LIMD1-Cry2 forming droplets after blue light exposure. Bottom: representative image of LIMD1 droplets formed at the same focal plane as FAs. Scale bar, 20 μm. **(e)** Left: FRAP analysis showing that LIMD1 droplets within FAs dynamically exchanges with their cytoplasmic counterparts. Right: quantification of FRAP analysis of the droplets indicated by the dashed box. LIMD1 fluorescence recover rapidly after photobleaching. Scale bar, 5 μm. **(f)** Representative images of Δ IDR-Cry2 showing few clusters and Δ LIM-Cry2 showing no clusters formed after blue light exposure. Scale bar, 20 μm. **(g)** Phase diagram of ΔLIM and LIMD1 proteins in different NaCl concentration. The red circles indicate phase separation, and the blue circles indicate no phase separation. **(h)** Representative images of LIMD1 phase separation at the FA sites with DMSO or Y27632 treatment. Scale bar, 20 μm. **(i)** Representative images showing that vinculin overexpression enhances LIMD1 droplets formation at the FA sites but not in the cytoplasm. Scale bar, 20 μm. **(j)** Representative images showing that vinculin-ActA relocates LIMD1 droplets to the mitochondria. Scale bar, 20 μm.

Since LIMD1 revealed force-dependent FA localization, we then interrogated whether mechanical force would affect LIMD1 phase separation. Cells expressing Opto-droplet-LIMD1 were treated with DMSO or Y27632 before being exposed to blue light. We observed that inhibiting the actomyosin contraction significantly reduces LIMD1 phase separation at the FA sites (Fig. 3h). Next, we asked how mechanical force would affect LIMD1 phase separation at the FAs. The interaction between vinculin and LIMD1 was detected by co-IP (Supplementary Fig. 3b). We then hypothesized that the conformational changes of vinculin induced by mechanical force^36^ may then locally increases LIMD1 concentration to allow phase separation to take place. In line with this, cells overexpressing vinculin greatly enhanced LIMD1 droplets formation at the FA sites while diminished droplets appearance in the cytoplasm (Fig. 3i). Furthermore, LIMD1 droplets relocated to the mitochondria when cells overexpressed the mitochondria-tethered vinculin (vinculin fused with ActA) (Fig. 3j and Supplementary Fig. 3c)^36, 37^. Collectively, these data reveal that LIMD1 could undergo phase separation both in vitro and in cells, and that its phase separation at the FAs is responsive to force.

Phase separation increases local concentration of certain protein components, driving non-membranous compartmentalization. We wondered whether the phase separation of LIMD1 would act semi-permeably to recruit specific FA proteins. Paxillin and zyxin were selected to represent early and late FA components. First, we asked whether LIMD1 could recruit paxillin or zyxin in vitro through phase separation. We performed in vitro co-phase separation experiments where both LIMD1 and paxillin or zyxin were present. Intriguingly, zyxin but not paxillin showed strong co-localization with LIMD1 in the phase-separated droplets (Fig. 4a and b). Of note, both paxillin and zyxin showed drastically reduced ability to undergo phase separation in vitro compared to LIMD1 (Supplementary Fig. 4a). Next, we explored whether the selective recruitment also occurred in cells. Opto-droplet-zyxin or Opto-droplet-paxillin showed minimal response to blue light exposure (Fig. 4c,d and Supplementary Video 15-16). However, by expressing both Opto-droplet-LIMD1 and GFP-tagged zyxin in the same cell, we detected nice and rapid co-localization of zyxin within LIMD1 droplets (Fig. 4e and Supplementary Video 17-18). On the contrary, recruitment of paxillin to LIMD1 droplets were barely detected when both Opto-droplet LIMD1 and GFP-paxillin were present in a cell (Fig. 4f and Supplementary Video 19-20). In addition, we also found the recruitment of late FA proteins Trip6 and VASP but not the early FA protein FAK to the LIMD1 Opto-droplets (Supplementary Fig. 4c-e and Supplementary Video 20-26). Taken together, our observations demonstrate that LIMD1 could recruit specific FA components through phase separation.

**Figure 4.**
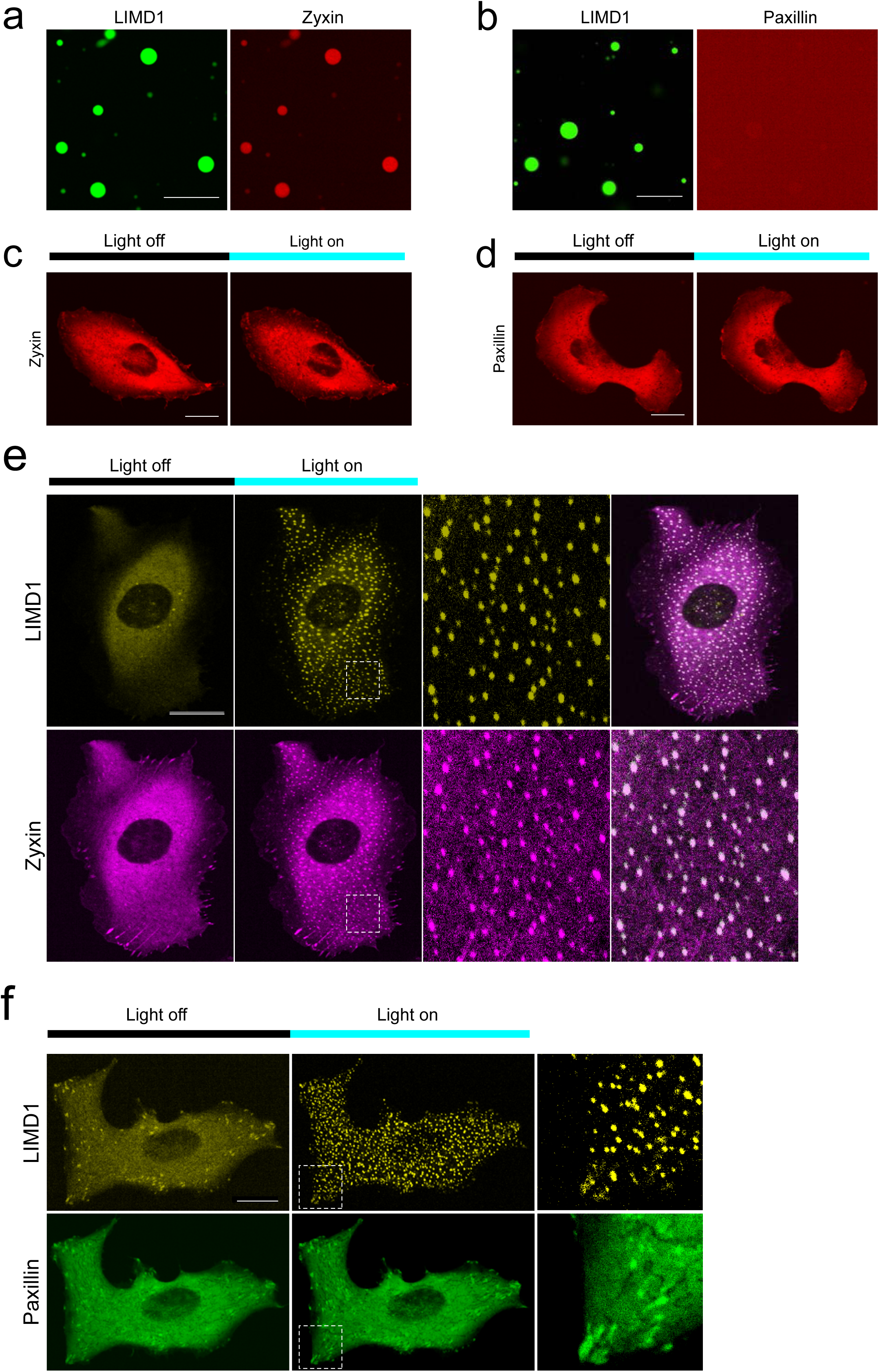
Phase separation of LIMD1 recruits specific focal adhesion proteins. **(a)** Representative fluorescence images showing that zyxin-mCherry protein shows strong co-localization with LIMD1-GFP protein in the phase-separated droplets. Scale bar, 20 μm. **(b)** Representative fluorescence images showing that paxillin-mCherry protein does not co-localize with LIMD1-GFP phase-separated droplets. Scale bar, 20 μm. **(c)** Representative images of zyxin-mCherry showing few clusters formed after blue light exposure. Scale bar, 20 μm. **(d)** Representative images of paxillin-mCherry showing no clusters formed after blue light exposure. Scale bar, 20 μm. **(e)** Representative images showing co-expression of LIMD1-Cry2 (yellow) and zyxin-GFP (magenta) in MDA-MB-231 cells forms multiple bright puncta containing both proteins after blue light exposure. Dashed boxes are zoomed-in on the right. Scale bar, 20 μm. **(f)** Representative images showing co-expression of LIMD1-Cry2 (yellow) and paxillin-GFP (green) in MDA-MB-231 cells. Dashed boxes are zoomed-in on the right. Scale bar, 20 μm.

Next, we asked whether perturbing LIMD1 phase separation would influence cellular mechano-response. Having observed that truncation of either the LIM domains (ΔLIM) or the IDR (ΔIDR) of LIMD1 drastically reduced its ability to phase separate, we then introduced these truncations back into LIMD1 KO cells, and interrogated the ability of these rescue cells to generate force. First, both of ΔLIM and ΔIDR were able to localize at the FAs (Supplementary Fig. 5a). Next, gel contractility assay and traction force microscopy were performed to evaluate cellular mechanical force. Interestingly, neither ΔLIM nor ΔIDR expressing cells were able to rescue defected gel deformation upon LIMD1 depletion (Fig. 5a and b). In contrast, reintroducing the full length LIMD1 back to these cells was able to recover cell contraction (Fig. 5a and b). Lastly, we compared the ability of these rescue cells to migrate directionally on a stiffness gradient gel. Notably, ΔLIM or ΔIDR expressing cells were still hampered in durotactic migration when compared with the LIMD1 KO cells rescued with the full-length LIMD1 (Fig. 5c and Supplementary Video 7-8, 27-29). Taken together, these results suggest that the phase separation property of LIMD1 may play an important role in regulating the mechanical responses of cells.

**Figure 5.**
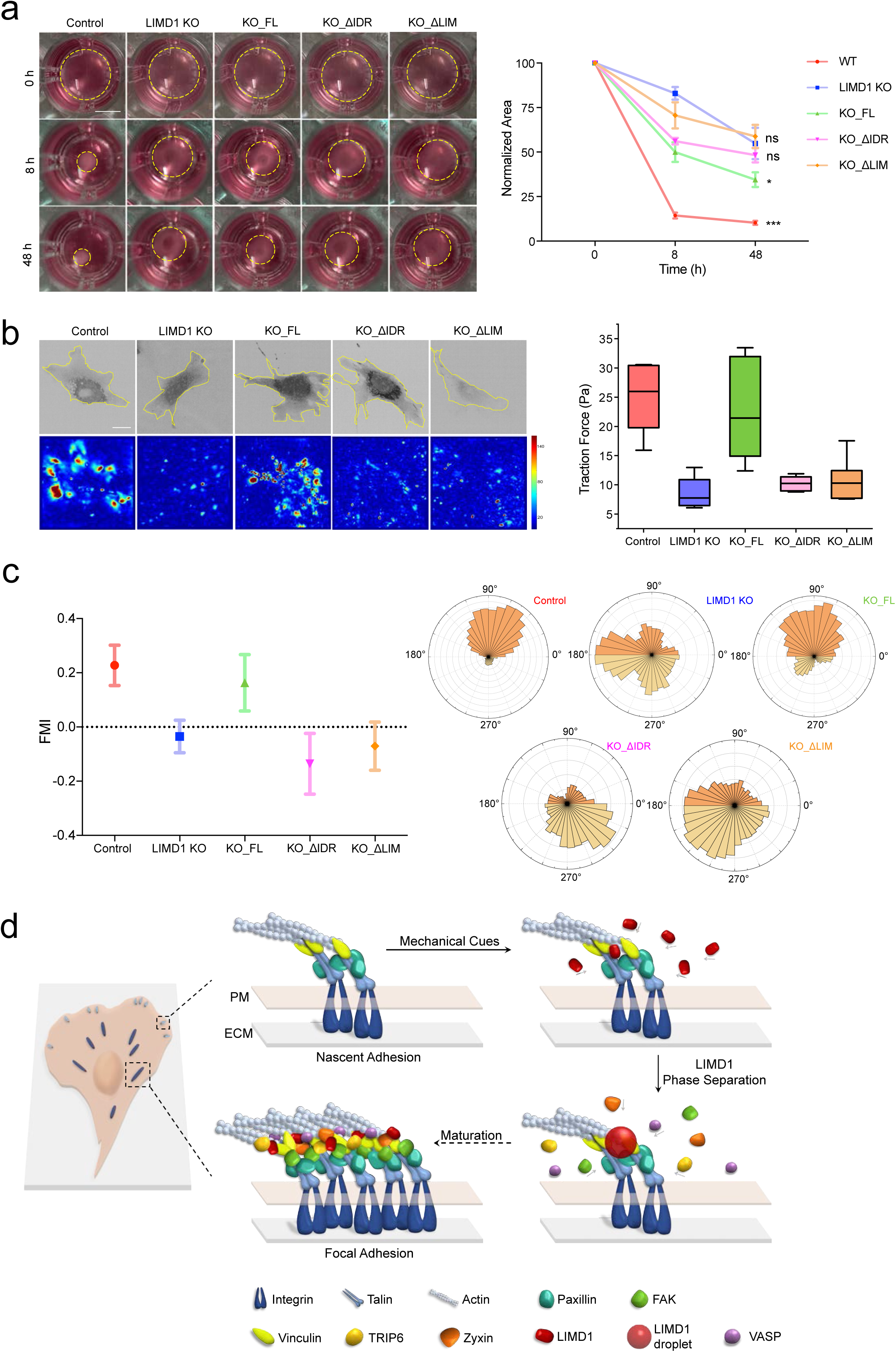
Perturbation of LIMD1 phase separation hinders cellular mechano-transduction. **(a)** Left: control, LIMD1 KO, KO_FL (LIMD1 KO rescued with full length LIMD1), KO_ ΔIDR (LIMD1 KO rescued with ΔIDR) and KO_ ΔLIM (LIMD1 KO rescued with ΔLIM) MEFs are cultured in 3D collagen gels. The morphology of cell-collagen gel mixture is recorded at 0 h, 8 h and 48 h. Scale bar, 5 mm. Right: quantification of collagen gel contraction using measurement of gel area (N = 3 experiments, error bars: 95% CI, *P < 0.05, ***P < 0.001 by Student’s *t* test for comparison of two groups). **(b)** Left: representative images of control, LIMD1 KO, KO_FL, KO_ ΔIDR and KO_ ΔLIM MEFs from traction force microscopy (TFM) experiments. Color spectrum indicates stress magnitude (Pa), with areas of low traction in blue and high traction in red. Right: quantification of total force from TFM experiments showing the different traction force of these five cell lines (n > 5 cells, error bars: 95% CI, *P < 0.05, **P < 0.01, and ***P < 0.001 by Student’s *t* test for comparison of two groups. ns, not significant). Scale bar, 20 μm. **(c)** Right: FMI of control, LIMD1 KO, KO_FL, KO_ ΔIDR and KO_ ΔLIM MEFs. N_Con_ = 55 cells, N_LIMD1 KO_ = 45 cells, N_KO_FL_ = 53 cells, N_KO_ ΔIDR_ = 34 cells, N_KO_ ΔLIM_ = 42 cells. Left: rose plots of angle distribution of five cell lines on the right. **(d)** Schematic model showing the LIMD1 phase separation at the FA sites triggered by mechanical cues in recruiting specific FA proteins during FA maturation.

In summary, our study has uncovered the role of LIMD1 in cellular mechanics and durotactic migration. We unveil the ability of LIMD1 to undergo phase separation and demonstrate that both the LIM domains and the IDR of LIMD1 synergistically contribute to its phase separation property. Furthermore, our data reveal the recruitment of specific late FA proteins by LIMD1 phase separation. We propose a model in which force-induced LIMD1 enrichment at the FAs and subsequent phase separation contribute to the recruitment of late FA proteins and ultimately facilitate cellular mechano-transduction and durotaxis (Fig. 5d). Phase separation reflects a demixing transition, in which a homogenous and well-mixed solution rearranges itself such that distinct regions of space are occupied by a specific concentration of species^38^. Compared to the hierarchical protein-protein interaction, LIMD1 phase separation may serve as a dynamic hub for rapid recruitment of specific proteins and may allow high permeability at the same time. It is an intriguing possibility that both LIM domain-dependent and IDR-driven interactions participate in achieving the specificity of protein recruitment. Phase separation is known to play roles in a variety of cellular processes, including formation of classical membrane-less organelles, signaling complexes, the cytoskeleton, and numerous other supramolecular assemblies^38^. The identification of FA protein phase separation illustrates another important physiological relevance of phase separation in response not only to biochemical inputs but also to biomechanical cues. Future studies to investigate the phase separation property of other FA components may extend our understandings in cellular crosstalk between sensation of the mechanical microenvironment and regulation of intercellular protein assemblies.

## METHODS

### Antibodies and reagents

The following antibodies were used in this study: anti-β-actin (ab8226) and anti-GAPDH (ab181602) were purchased from abcam; anti-α-tubulin (T9026) from Sigma-Aldrich; anti-GFP (598 and M048-3) from MBL; anti-paxillin (610620) from BD Biosciences; anti-LIMD1 (MABD85) from EMD Millipore; anti-vinculin (V9131) and anti-talin (SAB4200694) from Sigma-Aldrich; anti-P-FAK (Y397, 3283S) from Cell Signaling Technology; anti-RhoA (ARH04) from Cytoskeleton; anti-mouse (sc-2005) and anti-rabbit (sc-2004) horseradish peroxidase (HRP)-conjugated secondary antibodies from Santa Cruz Biotechnology. Donkey anti-Mouse IgG (H+L) Highly Cross-Adsorbed Secondary Antibody, Alexa Fluor 488 (A-21202) and 555 (A-31570) were purchased from Life Technologies.

Myosin II inhibitor (blebbistatin, 2946047) was purchased from EMD Millipore, Rho activator II (CN03) from Cytoskeleton, focal adhesion kinase-1 (FAK) inhibitor (PF 573228) from Sigma-Aldrich, ROCK Inhibitor (Y-27632, 13624S) from Cell Signaling Technology. Dimethyl sulfoxide (DMSO, 0247C211) was purchased from Life Science, Protease inhibitor mini tablets, EDTA-free (88666) from Thermo Fisher Scientific, Phosphatase inhibitor cocktail 3 (P0044) from Sigma-Aldrich. FluoShperes carboxylate-modified beads conjugated with 561 dye (0.1 μm, F8887) were purchased from Thermo Fisher Scientific. Protein A+G agarose beads (P2012) was purchased from Beyotime. CellTracker Green (C7025) and MitoTracker Red (M7512) were purchased from Invitrogen.

### Cell lines and cell culture

Mouse embryonic fibroblasts (MEFs) were a kind gift from James Bear laboratory at University of North Carolina at Chapel Hill. and human embryonic kidney 293T (HEK 293T) cells were kept by our laboratory. Human retinal pigment epithelium (hRPE) cells and MDA-MB-231 cells were generously provided by Yujie Sun laboratory (Peking University, Beijing, China).

Cells were cultured in Dulbecco’s modified Eagle medium (DMEM) supplemented with 10% fetal bovine serum (FBS; PAN, Biotech), 100 U/mL penicillin and 100 μg/mL streptomycin at 37°C with 5% CO_2_.

### Plasmids and transient infection

pcDNA3.1-CRY2-sfcherry plasmid was a kind gift from Yujie Sun laboratory. Human limd1, vinculin, zyxin, trip6 and vasp were cloned from cDNA library provided by Jiahuai Han laboratory (Xiamen University, Xiamen, China). Human paxillin was cloned from the HEK293FT cell-extracted cDNA library. Limd1, paxillin, vinculin, zyxin, trip6 and vasp were subcloned into lentiviral vector (pLVX-AcGFP-N1) or pLL5.0 vector for establishing stable cell lines.

HEK 293T, MEF, 231 and hRPE cells were transfected following the Neofect DNA transfection protocol (KS2000). After 24–48 h, cells were checked for fluorescent protein expression followed by imaging or harvesting cell lysates.

### CRSPR/Cas9-mediated LIMD1 gene knockout

The following single-guide RNAs (sgRNAs) and primers were used to generate LIM D1 knockout MEFs: mLIMD1-sgRNA-1: 5′-ATCGTGGTAGAGATCTCCCA-3′, mLIMD1-sgRNA-2: 5′-TCCTGTGTGAAATGCAGCAA-3′. A single MEF cell clone from lentivirus-infected pool cells was selected and verified by DNA sequencing.

### Western blotting

For western blotting, cells were washed with DPBS once and lysed in an appropriate volume of RIPA buffer (50 mM Tris-HCl, pH8.0, 150 mM NaCl, 1% Triton X-100, 0.5% Na-deoxycholate, 0.1% SDS, 1 mM EDTA and protease inhibitor cocktail) for 10 min on ice. Lysates were centrifuged at 13,572 g for 10 min and the supernatants were collected. Then, 5× SDS loading buffer was added to the supernatants and boiled for 10 min. Protein samples were run on 10% SDS–PAGE acrylamide gels and transferred onto nitrocellulose (NC) membranes by wet electrophoretic transfer, followed by primary and secondary antibody incubation at 4°C overnight or at room temperature for 2 h. The X-ray film was used to detect and record the band intensities. The fixed X-ray film was scanned and digital images were obtained. The band intensity was quantified by Fiji software (https://fiji.sc).

### Co-Immunoprecipitation

Cells were plated in 6 cm dishes and transfected with pLVX-AcGFP-N1 or vinculin-AcGFP and Limd1-mCherry, then were lysed by 350 μl of Pierce IP lysis buffer (25 mM Tris-HCl, 1 mM EDTA, 150 mM NaCl, 5% glycerol, 1% NP-40, pH = 7.4) for 15 min on ice. Lysates were centrifuged at 13,572 g, 4°C for 10 min and the supernatants were collected. 60μl of lysates were taken per sample and placed into a clean microfuge tube as “input” samples. Then 2.5 μg anti-GFP antibody was added into the remaining lysates and incubated on the rotating wheel at 4°C overnight. After that, 50 μl of protein A+G agarose beads were added to the lysates with antibody and incubated on the rotating wheel at 4°C for at least 3 h. Then the beads were collected and washed three times. 5 x sample loading buffer was used to resuspend the beads and “input” samples. Before separating by SDS-PAGE, the samples were boiled for 10 min and spin down briefly.

### Immunofluorescence and imaging analysis

Cells were plated on acid-washed coverslips coated with 10 μg/mL fibronectin overnight. Cells were then fixed with 4% paraformaldehyde at room temperature for 15 min, permeabilized in 0.5% Triton X-100 in PBS for 10 min, washed with PBS once for 5 min and blocked with 10% bovine serum albumin (BSA) for 1 h. Then the primary antibody was diluted 1:200 in 1% PBS and incubated for 1 h at room temperature. After washing with PBS three times, the coverslips were incubated with Alexa Fluor 488 or 555-conjugated secondary antibody for 1 h at room temperature. After another wash in PBS three times, the coverslips were mounted with Prolong Diamond Antifade with DAPI. Images were captured using Andor Dragonfly confocal imaging system.

### Live-cell imaging

Live-cell images were acquired with a 63×1.4 NA objective lens on a Andor Dragonfly confocal imaging system and EVOS M7000 imaging system (Thermo Fisher Scientific). Cells were plated on fibronectin (10 μg/mL)-coated glass-bottom cell culture dishes before imaging. Cells were maintained in DMEM supplemented with 10% FBS (PAN, Biotech), 100 U/mL penicillin and 100 μg/mL streptomycin at 37°C throughout the imaging process. Images were acquired at indicated intervals. For Opto-droplet phase separation imaging, cells were covered with tin foil to block light.

### Focal adhesion dynamics analysis

MEFs transiently transfected with mouse paxillin-GFP were placed in a heating chamber (37°C) on the stage of a time-lapse fluorescence microscope (IXplore TIRF; Olympus, Japan). Images were taken at 2.5-minute intervals and were processed for estimation of various parameters using Focal Adhesion Analysis Server (FAAS) (http://faas.bme.unc.edu). Focal adhesion longevity was calculated using FAAS and figures were plotted using Statistical Product and Service Solutions (SPSS).

### Fabrication of gradient and uniform polyacrylamide (PA) gels

Polyacrylamide gels with a stiffness gradient were generated as described before^1^ with mild modifications. A 65 μL drop of mixture containing 10% acrylamide, 0.5% bis-acrylamide and 2 mg/mL Irgacure (Sigma-Aldrich, 410896) was applied to glutaraldehyde-modified 24 mm glass coverslip, covered with a glass coverslip made hydrophobic by treatment with Repel-Silane. Gradients were generated by initially covering the acrylamide mix solution with an opaque mask and then slowly sliding it at a controlled speed while irradiating with a UV bench lamp. The mask was slid using an automatic syringe pump (Chemyx Fusion 200). To ensure complete polymerization, the whole acrylamide mix solution was first exposed to UV light for 10 min without covering (for the initial crosslinking), and then mask was slid at 40 μm/s for 10 min to produce the steep stiffness gradient gels. After gel photopolymerization, the hydrophobic glass coverslip was removed and the gel was washed with PBS thoroughly to remove unreacted reagents. The stiffness was measured with atomic force microscopy. To facilitate cell adhesion, fibronectin was covalently linked to the gels as described below.

### Functionalizing PA gel with Fibronectin

Polyacrylamide (PA) gels were made from 40% acrylamide and 2% bis-acrylamide mixed with 10% ammonium persulfate and 1% TEMED, where varying ratios of acrylamide and bis-acrylamide were used to create gels of known reproducible stiffness^2^. Gels were cast on glutaraldehyde-modified cover glasses. Premixed solutions (Soak solution: 137 mM NaCl, 5% (v/v) glycerol; 2x conjugation buffer: 0.2 M MES, 10% (v/v) glycerol, pH 4.5; 10x EDC: 150 mM, 29 mg/ml in DI water; 10x NHS: 250 mM, 29 mg/ml in DI water) were prepared. Soak solution was first pipetted onto each dish such that the gel was completely submerged (1.5mL), then dishes were incubated at room temperature for at least 1 h. After that, 1 part 10x EDC, 1 part 10x NHS, 3 parts DI water and 5 parts 2x conjugation buffer were mixed together. Removed all soak solution from dishes, then enough NHS/EDC solution was added to cover the gel surface (1.5mL) and dishes were incubated at room temperature for 30 min in the darkness. 20 μg/mL fibronectin was diluted in PBS (1:100), then removed the NHS/EDC solution. In the end, fibronectin solution was added to dishes, then dishes were incubated at room temperature for at least 35 min to allow for attachment of fibronectin. After that, PBS was added to each dish and dishes were stored at 4°C.

### Quantification and analysis of durotaxis on PA gel

Durotaxis was analyzed with a Andor Dragonfly confocal imaging system and EVOS M7000 imaging system using 10x objective. Cells were seeded onto the gradient PA gels and allowed to attach to gel surface for at least 8 h, then the samples were transferred to microscope culture chamber (37°C, 5% CO2). Time-lapse phase contrast images were taken every 15 minutes for 16 hours. Coordinates and distances of cell movement were tracked using the Fiji “Manual Tracking” plugin, then tracking data were imported into the “Chemotaxis Tool” plugin to generate statistic feature such as velocity, FMI (forward migration index), directionality and plot feature such as rose plot.

### Contractility assay

Cells were harvested and resuspended in desired medium at 1× 10^5^ cells/mL. 2% collagen lattice was prepared by mixing 10×PBS, NaOH and H_2_O in a tube on the ice, then collagen was added into the tube and followed by intensively mixed. 2% collagen was used to resuspend the cells. The mixture of cells and collagen were then seeded into a 48-well plate and cultured at 37°C 5% CO_2_. Photos of the collagen gels were taken at 8 h, 24 h and 48 h, and ImageJ was used to count the area of collagen gels at each time point, then a time-area curve was plotted.

### Fluorescence recovery after photobleaching (FRAP) assay

Cells were cultured in glass-bottom dishes at least 8 h before imaging. FRAP assay was performed on a ZEISS Laser Scanning Microscopes 880 at 37 °C. GFP signals in regions of interest (ROI) were bleached using a 488-nm laser beam at full power. The fluorescence intensity difference between pre-bleaching and the time point right after photobleaching pulse was normalized to 100%. The experimental control was to quantify fluorescence intensities of similar puncta/cytoplasm regions without photobleaching. In order to extract quantitative information from the FRAP curves, double-term fitting exponential equations were used in Microsoft Excel.

### Traction force microscopy (TFM)

Cells were stained with 1uM CellTracker Green for 50 min, and then plated on 4kPa polyacrylamide gels embedded with FluoShperes carboxylate-modified beads with 0.1-μm-diameter conjugated with 561 dye. The cells were allowed to adhere for at least 8 h. Cell spreading deformed the surfaces of the gels thus changing the position of the embedded beads. Fluorescence images of the embedded beads were captured on a Andor Dragonfly confocal imaging system. After taking the images of beads, trypsin/EDTA was added to detach the cells for at least 5 min and thus release forces mediated by cells on the gel. Subsequently, images of beads positions without cellular forces were obtained. The traction forces exerted by cells were analyzed by measuring the beads displacement induced by cells. Traction force analysis was performed using a MATLAB algorithm. The displacement of beads was calculated using a gradient-based digital image correlation technique (before and after trypsin/EDTA treatment). The corresponding gel deformation was obtained by two-dimensional Gaussian interpolation. The stress field (Pa) was calculated from the gel deformation by implementing a Fourier transform-based algorithm using the Boussinesq Green’s function.

Once traction force maps and fluorescent images were acquired, average traction force magnitude exerted by the whole cell can be calculated. Firstly, cell contours were extracted by built-in morphological functions in MATLAB. Then, cell contour was mapped onto traction force maps and set traction forces outside as zero. Finally, average traction force by the cell can be calculated by

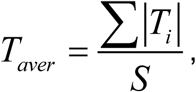

where, |*T*_*i*_ | indicates traction force magnitude per pixel and *S* represents the total area of cell contour.

### Prediction of disorder tendency

The disorder tendency of LIMD1 was predicted by IUPred (http://iupred.elte.hu).

### Constructs and Protein Expression

For recombinant expression in insect cells, DNA fragments encoding LIMD1 and ΔLIM were cloned into the pFastbac-HT B vector with a GFP tag and 6×His in the C terminus. Bacmids were generated using the Bac-to-Bac system (Invitrogen). Recombinant baculoviruses were generated and amplified using the sf9 insect cells, maintained in the SIM SF medium (Sino Biological Inc.).

For protein production, sf9 cells grown in the SIM SF medium (Sino Biological Inc.) were infected at a density of 3.0-4.0 × 106 cells/ml. 72 hours post infection, 0.5 liters of cell pellets were collected by centrifugation at 4000 g. Cell pellets were suspended in lysis buffer containing 20 mM Tris-HCl (pH8.0), 150 mM NaCl, 5 mM KCl, 10% glycerol, 20 mM imidazole, 1 mM tris (2-carboxyethyl) phosphine hydrochloride (TCEP), and protease inhibitor mixture (Complete EDTA-free, Roche), and were lysed by sonication and cleared by centrifugation (20,0 0 0 rpm). The proteins were purified with Ni-IDA (Smart-Lifesciences) chromatography, followed by size exclusion chromatography using Superdex200 Increase 10/300 (GE Healthcare) in buffer containing 20 mM Tris-HCl pH8.0, 500 mM NaCl. Finally, Proteins were concentrated and concentrations were determined with microspectrophotometry using the theoretical molar extinction coefficients at 280 nm. Protein purity was evaluated with Coomassie blue staining of SDS-PAGE gels.

### In vitro phase transition assay

#### Preparing coverslips and glass slides

Rinse glass slides and coverslips with ethanol twice, then dry with air. Wipe the slides and coverslips dry with lens cleaning tissue. Create an imaging chamber by attaching coverslip to the glass slide using double-sided tape placed parallel to each other, such that a rectangular chamber with two openings on opposite sides was made, named flow chamber.

#### Phase separation assays

The purified proteins LIMD1 and ΔLIM were diluted in a high salt solution containing 500 mM NaCl, 20 mM Tris (pH = 8.2), 1 mM DTT. To induce phase separation into liquid droplets at low salt conditions, proteins were diluted into the corresponding experimental buffers at a final concentration of 1.6 mM ATP, 4 mM MgCl_2_, 20 μM ZnCl_2_ and 1 mM DTT. Before use, the samples were thawed by incubating them at room temperature for 5 min. For imaging, droplets were observed by injecting protein-phase buffer mixtures into the homemade flow chamber for DIC or fluorescent imaging.

### Statistics and data display

The number of biological and technical replicates and the number of samples are indicated in figure legends, the main text and the Materials and Methods. Data are mean ± SEM or mean ± 95% CI. as indicated in the figure legends and Supplementary Figure legends. Student’s *t*-test and non-parametric test analysis were performed with GraphPad Prism 7.0. Data from image analysis was graphed using Prism and OriginPro 2019.

