## Supplementary Information for "LIMD1 phase separation contributes to cellular mechanics and durotaxis by regulating focal adhesion dynamics in response to force"

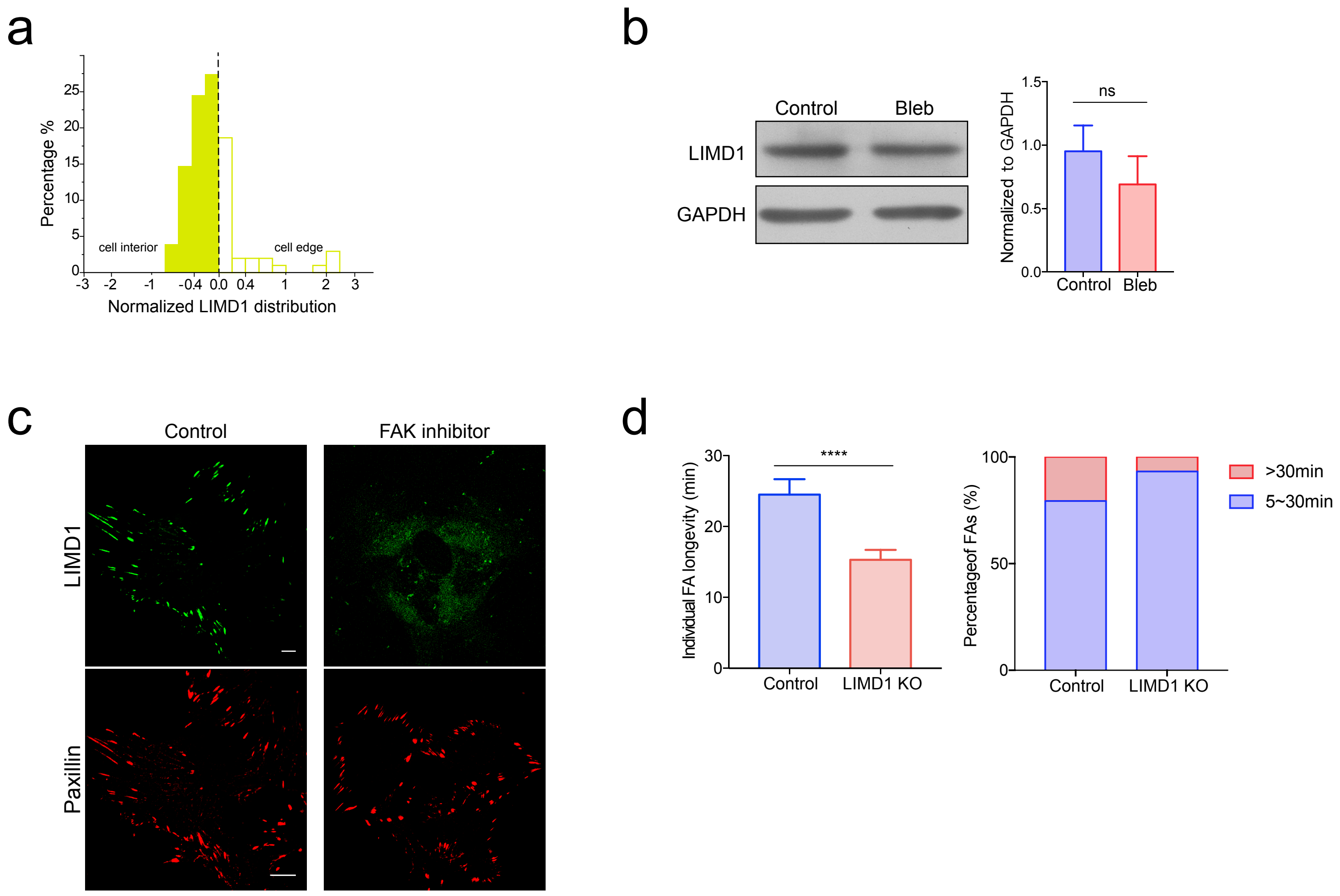


**Supplementary Figure 1**

(a) Quantification of LIMD1 distribution relative to paxillin within individual FAs. $N_{FA}$ = 102.

(b) Western blot showing the total protein level of LIMD1 in MEFs with or without blebbistatin treatment. GAPDH is used as loading control. Bar chart shows quantification of protein levels normalized to GAPDH in each condition (error bars: 95% CI, ns: not significant).

(c) Representative immunofluorescence images of LIMD1 and paxillin on stiff substrate with or without 10 μM FAK inhibitor treatment. Scale bars, 20 μm.

(d) Left: bar graphs showing individual FA longevity in control and LIMD1 KO MEFs (error bar: 95% CI, ***P < 0.001 by Student's t test for comparison of two groups). Right: bar graphs showing percentage of long-lived FAs.

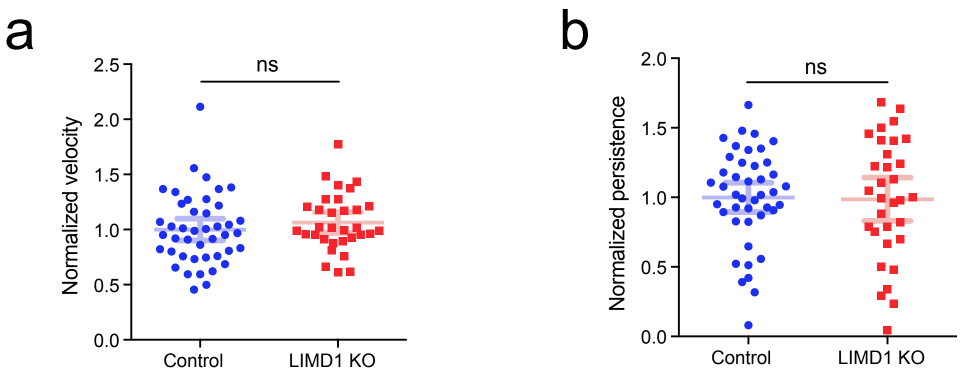


**Supplementary Figure 2**

(a) Quantification of velocity between control and LIMD1 KO MEFs (error bar: 95% CI, ns: not significant).

(b) Quantification of persistence between control and LIMD1 KO MEFs (error bar: 95% CI, ns: not significant).

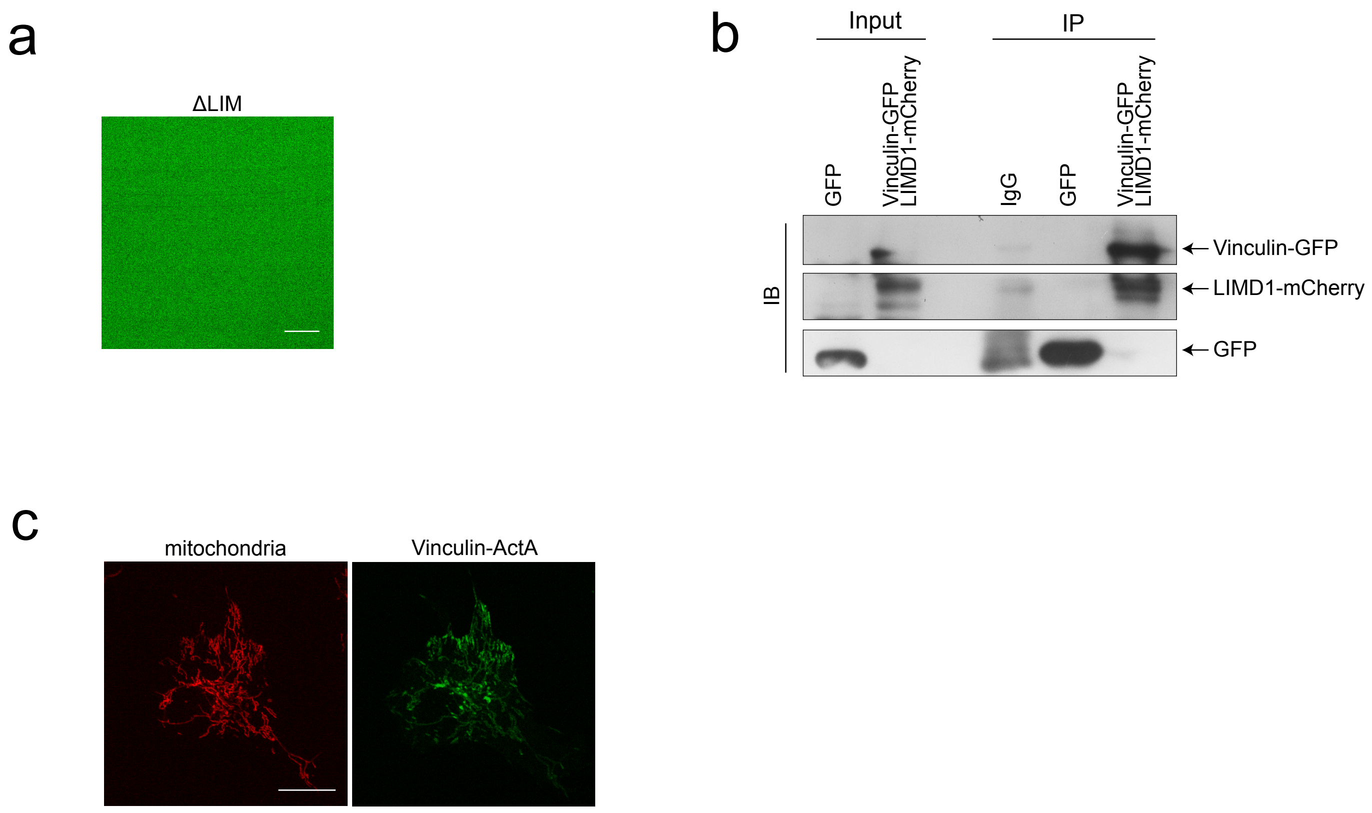


**Supplementary Figure 3**

(a) Representative images of ΔLIM protein tagged with GFP in phase separation buffer without formation of phase-separated droplets. Scale bar, 20 μm.

(b) Co-Immunoprecipitation assay showing interaction between vinculin-GFP and LIMD1-mCherry.

(c) Representative immunofluorescence images of mitochondria labeled with MitoTracker Red and vinculin-ActA. Scale bar, 20 μm.

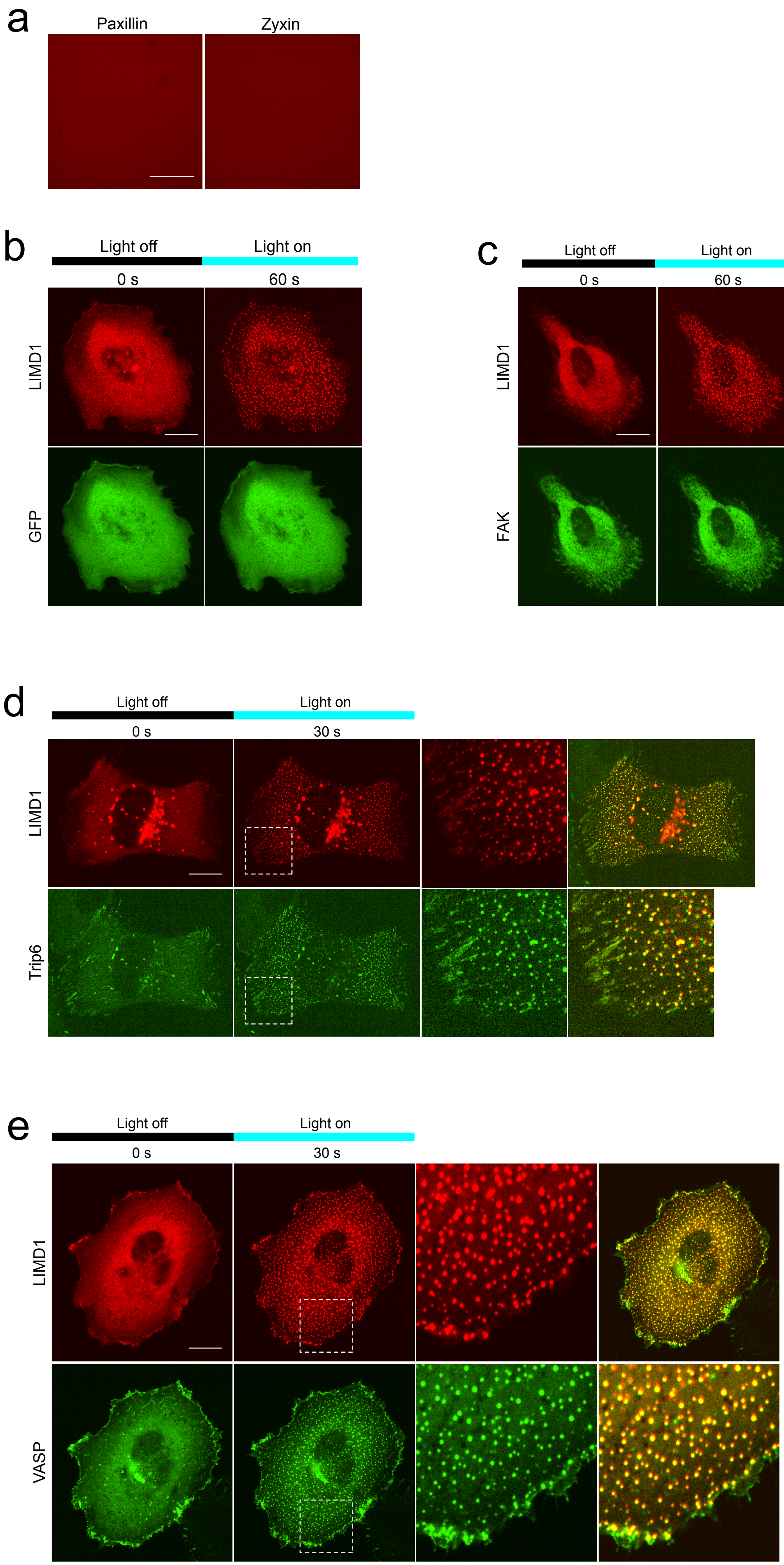
a
Paxillin
Zyxin
b
Light off
Light on
0 s
60 s
LIMD1
GFP
c
Light off
Light on
0 s
60 s
LIMD1
FAK
d
Light off
Light on
0 s
30 s
LIMD1
Trip6
e
Light off
Light on
0 s
30 s
LIMD1
VASP

**Supplementary Figure 4**

(a) Representative fluorescence images showing illin-mCherry and zyxin-mCherry proteins in phase separation buffer without formation of phase-separated droplets. Scale bar, 20 μm.

(b) Representative fluorescence images showing that GFP does t co-localize with LIMD1-Cry2 droplets in MDA-MB-231 cells after blue light exposure. Scale bar, 20 μm.

(c) Representative fluorescence images showing that FAK-GFP doe not co-localize with LIMD1-Cry2 droplets after blue light exposure. Scale bar, 20 μm.

(d) Representative fluorescence images showing that Trip6-GF xhibits strong co-localization with LIMD1-Cry2 droplets after blue light exposure. Scale bar, 20 μm.

(e) Representative fluorescence images showing that VASP-GFP hibits strong co-localization with LIMD1-Cry2 droplets after blue light exposure. Scale bar, 20 μm.

a

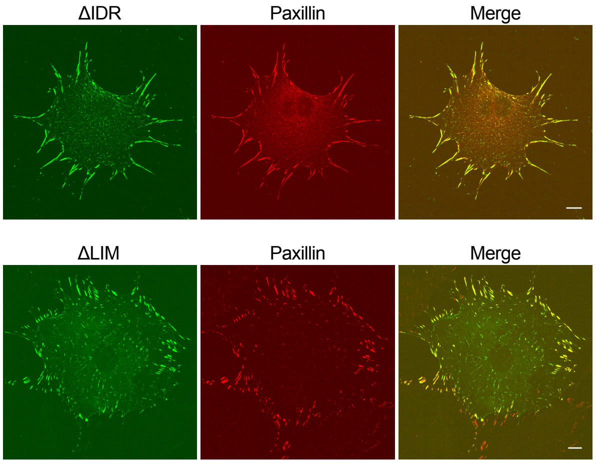


**Supplementary Figure 5**

(a) Representative images of ΔIDR-GFP and ΔLIM-GFP in cells. Scale bar, 20 μm.

**Supplementary Videos**

**Supplementary Video 1-2: Depletion of LIMD1 delayed cell spreading.** Time-lapse movies of WT (S1) or LIMD1 KO MEFs (S2). Time interval: 5 min, total time: 2 h. Scale bar: 20 µm.

**Supplementary Video 3-4: Delayed FA formation in LIMD1 KO cells upon washout of blebbistatin.** Time-lapse movies of WT (S3) or LIMD1 KO MEFs (S4) expressing vinculin-GFP under 40 µM blebbistatin for 1h and then washout with fresh medium. Time interval: 15 s, total time: 30 min. Scale bar: 20 µm.

**Supplementary Video 5-6: FAs dynamic analysis.** Live-cell TIRF was acquired for WT (S5) or LIMD1 KO MEFs (S6) expressing paxillin-GFP. Time interval: 150 s, total time: 90 min. Scale bar: 20 µm.

**Supplementary Video 7-8: Loss of durotaxis upon depletion of LIMD1 in cells.** Time-lapse movies of WT (S7) or LIMD1 KO MEFs (S8) migrating on stiffness gradient gels. Time interval: 15 min, total time: 16 h. Scale bar: 20 µm.

**Supplementary Video 9: Dynamic fusion events of LIMD1 droplets.** Time-lapse movie shows LIMD1 droplets fusing into larger ones. Time interval: 1 s, total time: 1 min. Scale bar: 4 µm.

**Supplementary Video 10: LIMD1 droplets showed rapid recovery upon photo-bleaching.** Time-lapse movie of LIMD1 droplet in a FRAP assay. Time interval: 30 s, total time: 30 min. Scale bar: 2 µm.

**Supplementary Video 11: Droplets formation of Opto-LIMD1 within cells.** Time-lapse movie of MBA-MD-231 cells expressing LIMD1-CRY2-sfCherry after blue light exposure. Time interval: 1 s, total time: 1 min. Scale bar: 20 µm.

**Supplementary Video 12: FA retained LIMD1 is dynamic.** Time-lapse movie of MEF cell expressing LIMD1-GFP in a FRAP assay, white arrowhead points to region of photobleaching. Time interval: 10 s, total time: 10 min. Scale bar: 5 µm.

**Supplementary Video 13-14: Truncation of the LIM domains or the IDR altered the LIMD1 droplet formation kinetics in cells.** Time-lapse movies of MBA-MD-231 cells expressing ΔLIM-CRY2-sfCherry (S13) or ΔIDR-CRY2-sfCherry (S14) after blue light exposure. Time interval: 1 s, total time: 1 min. Scale bar: 20 µm.

**Supplementary Video 15-16: Opto-zyxin or Opto-paxillin showed minimal response to blue light exposure.** Time-lapse movies of MBA-MD-231 cells expressing zyxin-CRY2-sfCherry (S15) or paxillin-CRY2-sfCherry (S16) after blue light exposure. Time interval: 1 s, total time: 1 min. Scale bar: 20 µm.

**Supplementary Video 17-18: Co-localization of zyxin within LIMD1 droplets.** Time-lapse movies of MBA-MD-231 cells expressing both LIMD1-CRY2-sfCherry (S17) and zyxin-GFP (S18) after blue light exposure. Time interval: 1 s, total time: 1 min. Scale bar: 20 µm.

**Supplementary Video 19-20: Co-localization detection of paxillin and LIMD1.** Time-lapse movies of MBA-MD-231 cells expressing both LIMD1-CRY2-sfCherry (S19) and paxillin-GFP (S20) after blue light exposure. Time interval: 1 s, total time: 1 min. Scale bar: 20 µm.

**Supplementary Video 21-22: Co-localization of TRIP6 within LIMD1 droplets.** Time-lapse movies of MBA-MD-231 cells expressing both LIMD1-CRY2-sfCherry (S21) and TRIP6-GFP (S22) after blue light exposure. Time interval: 1 s, total time: 1 min. Scale bar: 20 µm.

**Supplementary Video 23-24: Co-localization of VASP within LIMD1 droplets.** Time-lapse movies of MBA-MD-231 cells expressing both LIMD1-CRY2-sfCherry (S23) and VASP-GFP (S24) after blue light exposure. Time interval: 1 s, total time: 1 min. Scale bar: 20 µm.

**Supplementary Video 25-26: Co-localization detection of FAK and LIMD1.** Time-lapse movies of MBA-MD-231 cells expressing both LIMD1-CRY2-sfCherry (S25) and FAK-GFP (S26) after blue light exposure. Time interval: 1 s, total time: 1 min. Scale bar: 20 µm.

**Supplementary Video 27-29: Durotaxis detection of LIMD1 KO rescue cells.** Time-lapse movies of LIMD1 KO MEFs rescued with full length LIMD1 (S27), ΔLIM (S28) or ΔIDR (S29) respectively migrating on stiffness gradient gels. Time interval: 15 min, total time: 16 h. Scale bar: 20 µm.
